# Diet-Microbiome Analysis in a Healthy Cohort Reveals Potential Role of Intestinal Microbiota in Metabolism

**DOI:** 10.64898/2026.02.17.706416

**Authors:** Kenneth T Trang, Dalia Arafat Gulick, Jennifer Truell, Jiangze Tian, Rahul Bodkhe, Pranvera Hiseni, Kristin Gravdal, Tina Graceline Kirubakaran, Christina Casén, Rani Singh, Thomas R. Ziegler, Raylene A. Reimer, Colleen S. Kraft

## Abstract

Both preclinical and clinical studies have revealed the indisputable importance of intestinal bacterial community composition in pathogenesis of various disease states, from obesity to neurodegeneration. Diet remains one of the most important factors shaping human intestinal microbiota composition. In this study, we investigated diet-microbiome interactions in a healthy cohort of 88 participants from Atlanta and Calgary. We examine microbial composition (16S rRNA sequencing) with dietary records using Spearman Correlation tests with Benjamini-Hochberg multiple hypothesis correction to make community-level comparisons between dietary scores and microbial diversity index scores. Predictive models were used for molecular-level comparisons between microbial gene pathways and molecules. Among generalized dietary and microbial indices, we identified a negative association between dietary whole grain consumption and a microbial dysbiosis score. Comparisons between dietary food groups and bacterial family abundance reveal significant associations between dairy consumption and *Lactobacillaceae* abundance, dietary unsaturated to saturated fatty acid ratio and *Clostridia* Cluster Family XIII, salt intake and *Lachnospiraceae*, and consumption of ‘greens and beans’ and *Veillonellaceae*. Predictive models of microbial gene pathways and molecules reveal significant positive associations between several dietary fatty acids and microbial short-chain fatty acid fermentation pathways, and between dietary lignans and archaeal methanogenesis pathways. Overall, these associations may inform future explorations on specific dietary interventions to impact the gut microbiome.

**IMPORTANCE:** In this study, we compare dietary records and composition of intestinal microbes in a cohort of 88 participants. We identified associations between dietary consumption of dairy and the presence of dairy-consuming bacteria called *Lactobacteriaceae* and between consumption of dietary fats and the presence of fat-consuming bacteria called *Clostridia*. Using predictive analysis, we identify specific fatty acids associated with specific biochemical pathways found in *Clostridia* that might underlie these associations, in addition to an association between archaeal microbes and dietary consumption of estrogen-binding molecules called lignans, which are commonly found in whole grains and vegetables. Overall, our study generates useful associations between diet and intestinal microbes that can be tested in experiments that may help scientists use diet to control intestinal microbes in order to improve human health.

## INTRODUCTION

Diet is a modifiable risk factor and represents a major contributor to global morbidity and mortality burden^1,2^, with endogenous metabolism representing a highly complex biochemical interactions between humans and their external environment^3^. In addition to individual interactions within the food metabolome, host-associated factors such as genetics and intestinal microbiota add further complexity to these interactions. The latter represents hundreds of species with metabolic functions that bi-directionally interact with thousands of molecules to modulate disease pathogenesis^4,5^.

Previous investigations reveal numerous mechanisms underlying the role of intestinal microbiota metabolism on carbohydrates, including the microbial fermentation of dietary fiber into short-chain fatty acids (SCFAs). SCFA, including butyrate, propionate and acetate bind intestinal G protein-coupled receptors that inhibit histone deacetylase to reduce inflammation^6^. Lipid metabolism by intestinal microbiota is less understood, although several studies point towards the reduction of inflammation through the biotransformation of pro-inflammatory omega-6 and omega-3 polyunsaturated fatty acids (PUFAs) by microbes such as *Bifidobacterium* and *Lactobacillus* species^7^. High-fat diets may influence microbiome composition by increasing production of bile acids, which in addition to emulsifying fats, can also damage susceptible bacterial cell membranes^8^. Intestinal microbiota can transform primary bile acids into secondary bile acids, shifting the bile acid pool to impact survival and virulence of intestinal pathogens such as *Clostridium difficile*^9–11^ and *Vibrio cholera*^12^.

With this background, and the potential for precision microbiome engineering to modulate the role of diet in disease, in this study we investigate diet-microbiome interactions in the cohort of 88 healthy North American participants, comparing 16S rRNA microbiota sequencing reads in stool samples with dietary records using correlation tests with multiple hypothesis correction. We performed statistical comparisons including broad dietary and ecological index scores, a dysbiosis score, microbial family abundance, predicted molecules, and predictive microbial gene pathways. Our data reveal several significant family- and molecular-level associations that suggest mechanisms underlying the role of diet on microbiome composition in healthy adults.

## MATERIALS AND METHODS

### Study design

Participants were recruited from the communities surrounding the University of Calgary (Calgary, Canada) and Emory University Hospital (Atlanta, USA) [Fig. 1]. All participants provided written informed consent, and the study was approved by the Conjoint Health Research Ethics Board at the University of Calgary (REB18-0611) and the Emory University Institutional Review Board. To minimize the influence of any prevalent health conditions, healthy adult volunteers between the ages of 18 and 40 years were screened based on a health questionnaire, body mass index (BMI), blood pressure, dietary intake, and a fasted blood sample that was analyzed for lipids, fasting blood glucose, complete blood count (CBC), alanine aminotransferase (ALT), C-reactive protein (CRP), creatinine, and HbA1c.

**Figure 1:**
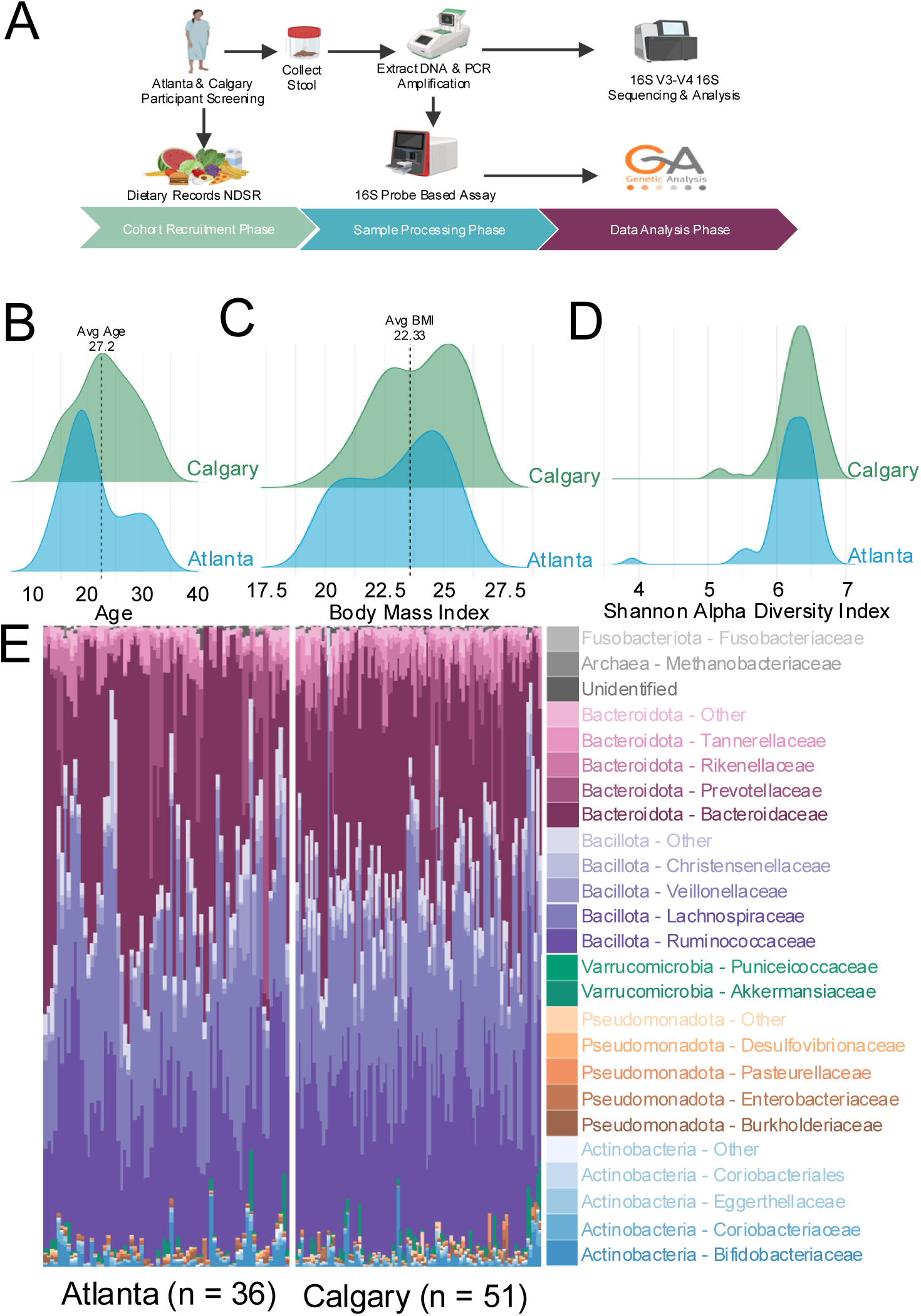
Demographic and 16S rRNA microbiota composition data from our North American cohort (n = 88). (A) After participants were recruited and screened, dietary records and stool samples were submitted, the latter of which underwent analysis via 16S rRNA sequencing and via commercial microbiome testing kit. (B) Distribution of age for Calgary and Atlanta cohorts. (C) Distribution of body-mass index for Calgary and Atlanta cohorts. (D) Distribution of Shannon Alpha Diversity index for Calgary and Atlanta cohorts. (E) 16S rRNA sequencing relative abundance results per sample.

To assess dietary intake, participants completed a 3-day diet record that included two weekdays and one weekend day. Dietary intake was analyzed with FoodWorks (Long Valley, NJ) using the Canadian or American Nutrient File as appropriate. Dietary intake was benchmarked against the Dietary Reference Intakes (DRIs) for each individual according to age and sex. Macronutrient intake (total fat, saturated fat, protein, carbohydrates and sugar) were within the *Acceptable Macronutrient Distribution Ranges* for inclusion.

Based on dietary records, we used Nutrition Data System for Research (NDSR)^13^ to calculate the Healthy Eating Index 2015 (HEI-2015) score^14^, an assessment tool that calculates individual component scores for each evaluated food group, with total scores ranging from 0-100 and higher values indicating greater alignment with United States Department of Agriculture Food and Nutrition Service Dietary Guidelines for Americans. We also used NDSR to predict molecules from dietary records.

Participants also completed a demographics questionnaire and Godin’s Leisure Time Exercise questionnaire^15^. The inclusion and exclusion criteria were:

<u>Inclusion Criteria</u>: Healthy male and female subjects who:

1. Were not overweight or obese (BMI ≥ 18.5 kg/m^2^ and ≤ 24.9 kg/m^2^)
2. Were between 18 and 40 years of age
3. Had regular bowel movements (no diarrhea or constipation according to Bristol stool chart)
4. Had maintained a stable body weight (within 3 kg) for at least 3 months before enrollment

<u>Exclusion Criteria</u>: Subjects with:

1. Chronic disease including but not limited to intestinal disease (e.g., Crohn’s disease, ulcerative colitis, irritable bowel syndrome, persistent or infectious diarrhea, chronic constipation, esophageal reflux), type 1 or 2 diabetes, cardiovascular disease, dyslipidemia, depression or anxiety, cancer, liver or pancreas disease
2. Major gastrointestinal surgeries (excluding appendectomy)
3. Pregnant or lactating
4. Used tobacco
5. Had taken antibiotics/antifungals/antivirals in the preceding 3 months
6. Currently consume probiotic or prebiotics in supplement form (note that foods containing prebiotics and/or probiotics such as probiotic yogurt or a granola bar with prebiotic in it would be allowed unless consumed in high doses)
7. Taking laxatives, proton pump inhibitors or over the counter anti-diarrheal medication
8. Investigational drug or vaccine
9. Following a diet or exercise regimen designed for weight loss
10. Had a BMI greater than 24.9 or less than 18.5 kg/m^2^
11. Consumed more than 2 standard alcoholic drinks per day in males, consume more than 1 drinks per day in females

### Stool Processing and GA-map® Dysbiosis Test Lx

After screening, each patient was provided a stool kit containing a pair of gloves, stool collection frame, a stool collection tub, a sterile specimen container, a sterile wooden scraper, a label, small biohazard bag, ice packs with an instruction sheet. Two stool samples were collected at home by participants and stored in their home freezer (-20°C) until delivered to the investigators collection site where they will be stored at -80°C. (ideally within 1-3 days).

A total of 176 stool samples were sent to Genetic Analysis AS (Oslo, Norway) and processed for the GA-map® Dysbiosis Test Lx^16^, a commercial test that measures bacterial abundance using a predetermined set of 48 magnetically coupled DNA probes complementary to 16S rRNA gene sequences in order to predict dysbiosis severity and functional profiles. To process samples, fecal samples were homogenized before undergoing mechanical and chemical disruption to isolate bacterial DNA. Following PCR amplification of extracted DNA using primers specific for V3-V9 16S regions, amplicons were hybridized to DNA probes, with fluorescence measured by Luminex® 200TM instrument (Luminex Corporation). Results were then analyzed by Genetic Analysis AS in reference to healthy cohort in order to calculate a Dysbiotic Index (DI): with values DI = 1-2 indicating normobiosis, DI = 3 indicating mild dysbiosis, and DI = 4-5 indicating severe dysbiosis^16^.

### 16S rRNA sequencing and analysis

To assess the variation in microbiota composition within our healthy cohort, we characterized the gut microbiota composition of participants using V3-V4 16S sequencing of two stool samples per volunteer collected one week apart. DNA from 176 stool samples was extracted by Genetic Analysis and shipped to Emory University’s Integrated Genomics Core for V3-V4 16S rRNA Illumina MiSeq 2x300 paired-end sequencing using 341F and 805R primers. DADA2 was used to perform quality control of raw reads and assign amplicon sequence variants (ASVs), with taxonomic assignment using SILVA v132 database^17^. Further statistical analysis and visualization were performed with the following R packages: phyloseq^18^, microshades^19^, phangorn^18^, and ggplot2^20^. To visualize and quantify differences in our microbiota data, we utilized Principal Component Analysis with weighted UniFrac distances (PCoA-UniFrac) to project our hyperdimensional microbiota data along two composite axes, using PERMANOVA for statistical comparisons between groups. To compare the diversity of each microbiota sample, we calculate a Shannon diversity index score, an alpha diversity metric that considers both species richness and evenness, in addition to using the dysbiosis index score from the commercial test described above. Predicted functional microbial gene pathway analysis was performed with PICRUSt2^21^. All comparisons between diet and 16S rRNA microbiota data were performed with stool collected from Week 1.

## RESULTS

### Patient demographics

We recruited 88 young (age: range = 18-38, mean = 27.4, SD = 5.5), adult participants with healthy body mass index (BMI: range = 18.4-25, mean = 22.3, SD = 1.9) [Fig. 1A, B, C]. 36 participants were from Atlanta, Georgia in the USA and 51 participants were from Calgary, Alberta in Canada, with no significant difference between age (P = 0.108; two-sided t-test) [Fig. 1B] and only slightly lower BMI found in Atlanta compared to Calgary (P = 0.027; uncorrected two-sided t-test) [Fig. 1C]. Overall, we conclude that our cohort represented a healthy group of adult participants that minimizes demographic and disease-related confounding factors.

### 16S stool microbiota of healthy participants in Atlanta and Calgary

To understand potential confounding relationships between diet and microbiota composition, we compared 16S rRNA microbiota composition with demographic factors. We observed no significant differences between 16S rRNA microbiota composition in samples collected during the first and second week using either Shannon Diversity Index (P = 0.7923; two-sided t-test) [Fig. S1A] or PCoA-UniFrac (P= 0.432; PERMANOVA) [Fig. S1B], suggesting that gut microbiota composition does not vary significantly within one week in our cohorts. Shannon alpha diversity did not differ between Atlanta and Calgary (P = 0.1338; two-sided t-test) [Fig. 1D], with only a marginally significant difference between sites when plotted on PCoA-UniFrac axes (P = 0.021; PERMANOVA) [Fig. S1E]. Although Shannon diversity was not correlated with BMI (P = 0.7876; Spearman’s Correlation) [Fig. S1C], we observed only a marginal negative relationship between age and Shannon Diversity (ρ = -0.17; P = 0.023; Spearman’s Correlation) [Fig. S1D]. In addition, we observed no significant relationship between 16S rRNA gut microbiota composition PCoA-UniFrac with either BMI (P = 0.122; PERMANOVA) [Fig. S1E] or age (P = 0.105; PERMANOVA) [Fig. S1F].

Across all samples, Bacillota (Relative abundance: mean = 56.2%, SD = 11.1%) and Bacteroidota (Relative abundance: mean = 39.5%, SD = 12.3%) were the two phyla that accounted for the largest percentage of identified sequence reads per participant, with Actinomycetota, Proteobacteria, and Verrucomicrobia the subsequent next most abundant phyla [Fig. 1E]—largely reflecting 16S rRNA microbiota phyla abundances for Westerners reported by The Human Microbiome Project^22^. We therefore conclude that our microbiota data reflects our current understanding of what constitutes the microbiota composition of a healthy younger adults, with negligible confounding factors of sample timing, geography, age, and BMI.

### Dietary records of healthy patients in Atlanta and Calgary

In our full cohort, the mean total HEI-2015 score was within the range of “needs improvement” (mean = 68.0, SD = 10.6), although notably higher than the mean total HEI-2015 score previously reported in the USA (HEI = 56.6) [Fig. 3A], with no significant difference (P = 0.22; t-test) between total HEI-2015 scores between Calgary and Atlanta [Fig. 3A]. We additionally observed no significant relationships between HEI-2015-Total and age (P = 0.105) or BMI (P = 0.89). To explore further confounding factors, we performed a Spearman’s correlation test between 207 NDSR-identified nutrients and demographic factors (age and BMI) [Table S1].

Although all other associations were not significant after multiple hypothesis correction (FDR > 0.2), there were negative associations between predicted dietary glycemic index (both glucose and bread reference) and age (P < 0.0001) [Table S1, Fig. S1F]. Overall, we conclude that although our cohort features a wide variation in HEI scores, our participants on average have a healthier diet than the average USA population, with minimal confounding demographic factors.

### Associations between dietary food groups and fecal 16S rRNA microbiota in healthy participants

To access the relationship between dietary intake and microbiota composition within a healthy cohort, we compared patterns in HEI-2015 total and food groups with our 16S rRNA microbiota data from Week 1. PCoA-UniFrac analysis revealed no significant relationship between 16S rRNA gut microbiota composition and total HEI-2015 score (P = 0.097; PERMANOVA) [Fig. 2B]. We next performed a Spearman’s correlation test with BH correction between the 14 HEI-2015 (total or component groups) scores, and the Shannon Diversity Index or GA-map® Dysbiosis Test Lx Dysbiotic Index (DI) scores [Table S2]. We observed a significant negative correlation between DI and ‘HEI-2015 Whole Grain’ component score (P = 2.8x10^-3^; FDR < 0.2) [Fig. 2D], consistent with findings from a previous randomized control intervention demonstrating that whole grain consumption reduces dysbiosis^23^.

**Figure 2:**
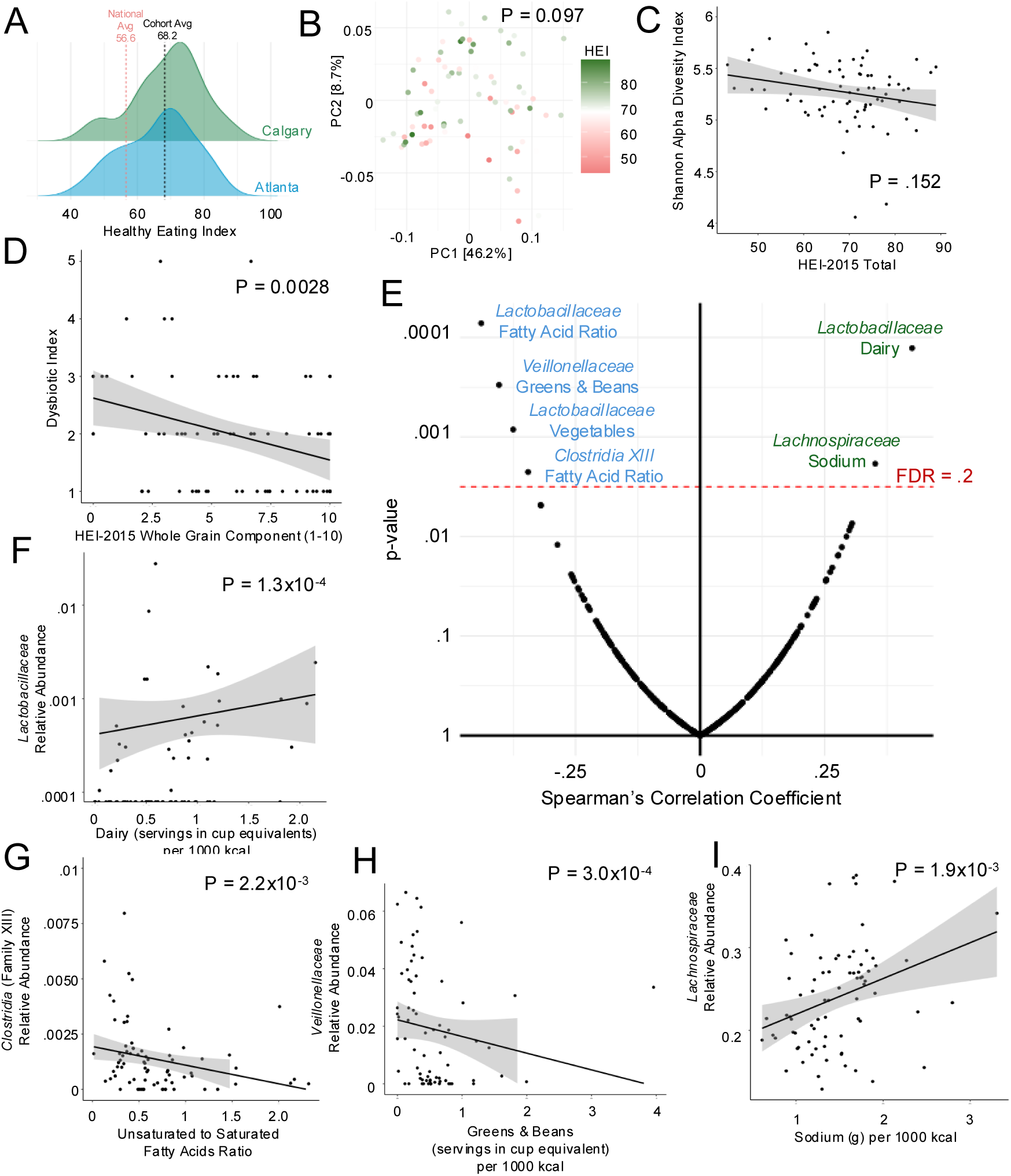
Dietary records compared to microbial family relative abundances. (A) Distribution of healthy eating index scores for Calgary and Atlanta cohorts. (B) Principal Component Analysis (PCoA) of 16S rRNA microbiota composition with green-red shading indicating HEI levels per sample. (C) Spearman correlation analysis between ‘HEI-2015 total’ and Shannon Alpha Diversity index. (D) Spearman correlation analysis between ‘HEI-2015 Whole Grain Component’ and Dysbiotic index. (E) Spearman’s correlation test with Benjamini-Hochberg multiple hypothesis correction between the 13 measured dietary groups and the 16S rRNA microbiota composition relative abundance of the thirty most represented bacterial families. Plotted is Spearman Correlation Coefficient against p-value, with red dotted line indicating false discovery rate = 0.2. Labels are included for the six comparison that are significant after BH correction. (F) Spearman correlation analysis between consumption of ‘Dairy (servings in cup equivalent) per 1000 kcal’ and relative abundance of Lactobacillaceae. (G) Spearman correlation analysis between ‘Unsaturated to Saturated Fatty Acid Ratio’ and relative abundance of Clostridia Family XIII. (H) Spearman correlation analysis between consumption of ‘Green & Beans (servings in cup equivalent) per 1000 kcal’ and relative abundance of Veillonellaceae. (I) Spearman correlation analysis between consumption of ‘Sodium (g) per 1000 kcal’ and relative abundance of Lachnospiraceae.

To explore more precise relationships between diet and microbiota composition, we performed pairwise Spearman’s correlation test with BH correction between the 13 measured dietary groups and the 16S rRNA microbiota composition relative abundance of the thirty most represented bacterial families [Table S3], resulting in 6 out of 390 (1.54%) significant correlations (FDR < 0.2 [Fig. 2E]). We observed a significant positive Spearman’s correlation between the consumption of dairy and the relative abundance of *Lactobacillaceae*, a family of gram-positive bacteria used to ferment milk into yogurt and cheese (P = 1.3x10^-4^; FDR < 0.2) [Fig. 2F]. There were also significant negative Spearman’s correlations between *Lactobacillaceae* and the dietary unsaturated to saturated fatty acid ratio (P = 7.21x10^-5^; FDR < 0.2) and with total vegetable intake (P = 8.4x10^-4^; FDR < 0.2) [Fig. 2E]. Similarly, we observed a negative correlation between the relative abundance of anerobic fermentating *Clostridia* Cluster Family XIII and the unsaturated to saturated fatty acid ratio (P = 2.2x10^-3^; FDR < 0.2) [Fig. 2G].

There was a significant negative association between the dietary intake category ‘Greens and Beans’ and the *Veillonellaceae* (P = 3.0x10^-4^; FDR < 0.2) [Fig. 2H], a less well-studied family of gram-negative anaerobic bacteria associated with green leafy vegetables, the main source of dietary nitrate that *Veillonellaceae* characteristically use for anaerobic respiration^24–26^. In addition, we observed a positive Spearman’s correlation between the relative abundance of *Lachnospiraceae* and consumption of sodium (P = 0.0019; FDR < 0.2) [Fig. 2I], notable as several preclinical studies report that dietary intake of salt increases *Lachnospiraceae* abundance^27–29^. Overall, we identify several diet-microbiota associations in our cohort that reflect mechanisms previously found in preclinical models and clinical studies.

### Associations between dietary nutrients and predicted microbial metabolism from fecal 16S rRNA sequencing in healthy participants

To investigate relationships between molecules and intestinal microbiota function, we performed a Spearman’s correlation coefficient test with multiple hypothesis testing correction between 364 microbial metabolic pathways predicted by PICRUSt2 and 224 additional factors, including HEI-2015 total and component scores, predicted dietary molecules, demographic factors, and DI [Table S6]. Among the 81,536 comparisons made, 96 associations (0.12%) were significant after multiple hypothesis correction (FDR < 0.2) which included 37 predicted metabolic pathways and 35 tested covariates [Figure 3J]. HEI-2015 total score was negatively associated with formaldehyde assimilation pathway (P = 2.1x10^-4^) [Fig. 3A]. Additionally, the Dysbiotic Index was significantly associated with the following three microbial metabolic pathways: *Enterococcus*-associated Peptidoglycan Synthesis (IV) (P = 1.3x10^-4^) [Fig. 3B], Archaeal-associated Glycolysis (V) (P = 1.1x10^-4^) [Figure 3J], and *Bifidobacterium*-associated Sucrose degradation (P = 1.9x10^-4^) [Figure 3J]. Although the mechanisms underlying these associations is unclear, *Enterococcus* peptidoglycan stimulates inflammation^30,31^, which may underlie the inflammatory symptoms characteristic of patients with dybiosis^32^.

**Figure 3:**
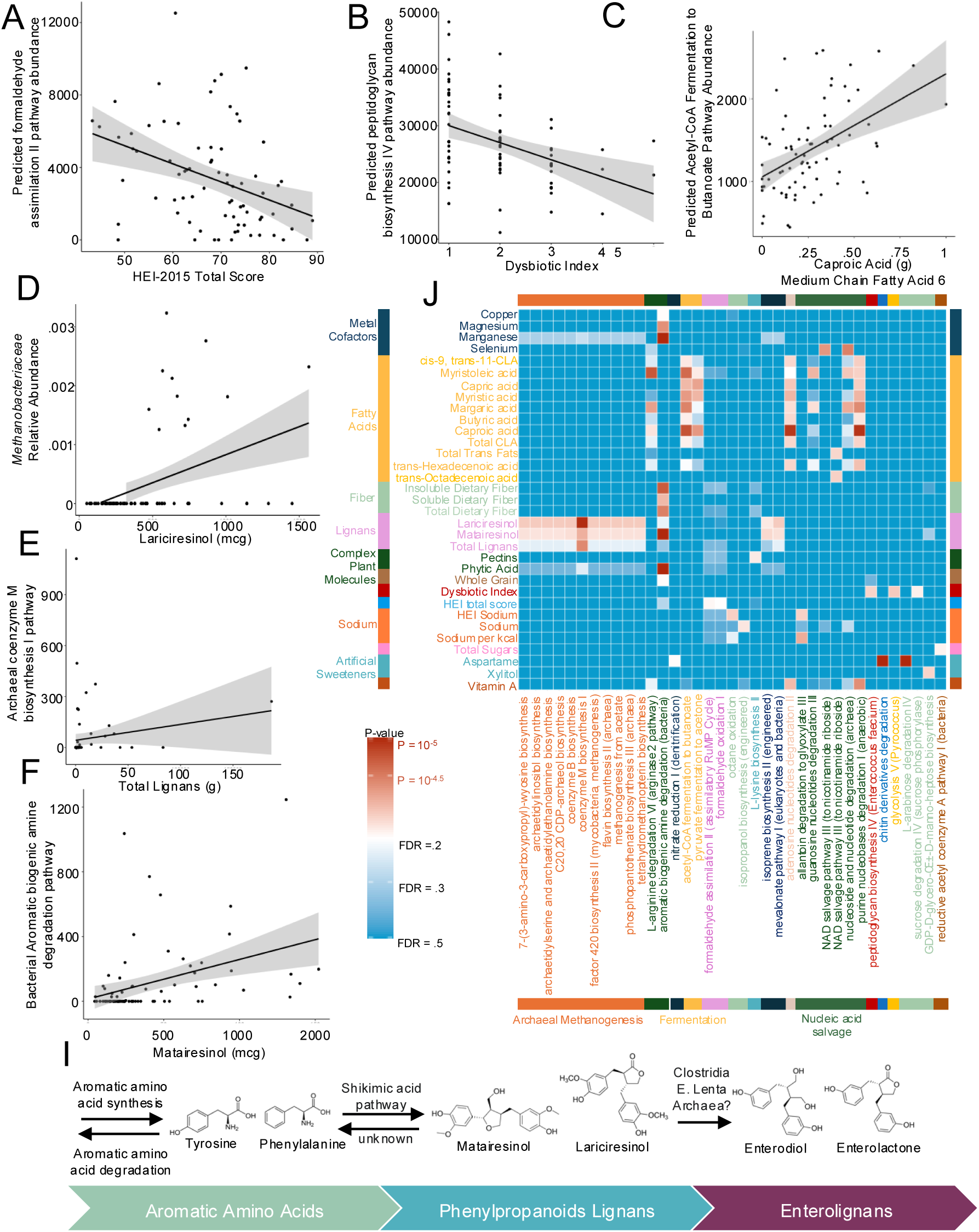
Predicated molecules and additional factors compared with predicated microbial functional pathways. (A) Spearman correlation analysis between ‘HEI-2015 Total score’ and ‘Formaldehyde assimilation II pathway’. (B) Spearman correlation analysis between ‘Dysbiotic index’ and ‘Peptidoglycan biosynthesis IV pathway’. (C) Spearman correlation analysis between ‘Caproic Acid (g) Medium Chain Fatty Acid 6’ and ‘Acetyl-CoA fermentation to butanoate pathway’. (D) Spearman correlation analysis between Lariciresinol (mcg) and relative abundance of Methanobacteriaceae. (E) Spearman correlation analysis between Total lignans (g) and ‘Archaeal coenzyme M biosynthesis I pathway’. (F) Spearman correlation analysis between ‘Matairesinol (mcg)’ and ‘Bacterial Aromatic biogenic amine degradation pathway’. (I) Lignan metabolic pathway depicting the conversion of lignans Matairesinol and Lariciresinol to enterolignans or aromatic amino acid precursors. (J) Heatmap of the 96 significantly associations between predicated molecules with additional factors and predicated microbial functional pathways.

Underscoring the functional congruence of our significant associations, all 12 significant associations (FDR < 0.2) involving *Clostridia*-associated microbial fermentation pathways (either ‘acetyl-CoA fermentation to butyrate’ or ‘pyruvate fermentation to acetone’) were positively associated with dietary fatty acids [Fig. 3J], which through upstream fatty acid oxidation would produce the acetyl-CoA, utilized in these two fermentation reactions^7^. In reference to the association reported above between *Clostridia* Cluster Family XIII and dietary fatty acids, these significant positive associations serve as a potential elaboration of the metabolic abilities unique to *Clostridia* that promote survival and replication induced by ingestion of dietary fatty acids.

Of 23 of the 96 significant associations involved Archaeal methanogenic-associated metabolic pathways, all were positively correlated with lignans (matairesinol, lariciresinol, or total lignans), plant-derived secondary metabolites molecules found commonly in whole grains and seeds [Fig. 3J]. Lignans are synthesized from aromatic amino acids and converted by intestinal microbiota into enterolignans, metabolites associated with estrogen-associated carcinogenesis due to their ability to bind to estrogen receptors^33–35^. We found a positive association between matairesinol and bacterially-associated aromatic amino acid degradation (P = 4.4x10^-6^; Spearman’s correlation) [Fig. 3F], suggesting matairesinol may be degraded into precursor aromatic amino acids within the gut, which may induce aromatic amino acid degradation pathways in intestinal bacteria. Aromatic amino acid degradation was not significantly associated with any of the three dietary aromatic amino acids (FDR > 0.2), but there were significant positive associations (FDR < 0.2) with matairesinol, plant-associated molecules (phytic acid, total dietary fiber, soluble dietary fiber, and whole grains), and metal cofactors (copper, manganese, and magnesium). Although speculative, this network of correlations may suggest a potential cascade of transkingdom microbial metabolism within the gut lumen induced by dietary intake of lignans.

## DISCUSSION

This study examined a small cohort of healthy, young North American adults to explore diet-microbiota interactions. We did not identify significant associations between generalized dietary and microbial indices such as HEI and alpha diversity, as in a larger previous study^36^; however, our study showed a negative association with a Dysbiosis index and consumption of whole grains, recapitulating results from a randomized intervention demonstrating that whole grain consumption can reduce dysbiosis and plasma IL-6^23^ and highlighting the utility of a functional dysbiotic index over a diversity index to describe microbiota composition.

We identified six significant associations between bacterial families and dietary food groups that reflect both clinical studies and well-established preclinical mechanism underlying microbiota metabolism of molecules. Previous studies have identified dairy consumption as a major dietary factor modulating intestinal microbiota composition^37^, with some large cohort studies reporting increases in *Lactobacillus* species^38,39^ while others report increases in *Bifidobacterium* species^40,41^. In our modest sized cohort, we identified only a positive significant association with *Lactobacillaceae*. In fact, *Bifidobacterium* and dairy consumption were not associated at all in our cohort (ρ = 0.014; P = 0.75; Spearman Correlation), suggesting that other poorly understood factors may underlie the competition between *Lactobacillaceae* and *Bifidobacterium* competition for dairy metabolites.

We additionally identified an association between *Clostridia* XII and dietary fatty acid ratio, with predictive analysis suggesting that this relationship is associated with specific dietary fatty acids and *Clostridia* butyrate fermentation pathways. While many studies have demonstrated mechanisms underlying the role of high fat diets on *Clostridioides difficile* infection (CDI) ^42–45^, *Clostridia* include a large class of commensal and beneficial bacteria. Many preclinical studies suggest mechanisms underlying the role of *Clostridia* species to ameliorate disease in high-fat diet contexts and inhibit CDI ^46–48^. However, the therapeutic role of *Clostridia* in high-fat dietary contexts remains largely unexplored in human contexts.

Our study also identified a significant negative association with ‘Greens and Beans’ and *Veillonellaceae*, a poorly characterized family of anerobic bacteria capable of nitrate respiration and associated with CDI in Crohn’s disease patients^24^. Although *Veillonellaceae* may convert nitrate to nitrite for energy production, green leafy vegetables contain both high levels of nitrate and nitrite^49^, the latter of which inhibits *Veillonellaceae* nitrate reduction through negative feedback. Several oral microbiome intervention studies demonstrate that supplemental dietary nitrates lead to an increase in aerobic nitrate respirators *Rothia* and *Streptococcus* and a decrease in anerobic nitrate respirators *Veillonella* and *Prevetolla* within the oral environment^50–52^.

Although another study reported a weakly significant positive association between a pro-inflammatory diet poor in vegetables, and *Veillonella rogosae* abundance in stool samples from IBD patients^25^, our study provides a clear negative association between dietary consumption of ‘Green and Beans’ and *Veillonellaceae* abundance. This provides a foundation for future studies exploring the potential role of dietary nitrates, nitrites, and *Veillonellaceae* nitrate respiration in the metabolism of plant-based diets and resistance to CDI.

While almost all significant associations identified in this cohort reflect well-established mechanisms and clinical associations, we found a significant positive association between archaeal methanogenesis and two lignans: matairesinol and lariciresinol. Currently, only one well-characterized pathway has been identified to convert lignans to enterolignans, involving a multistep pathway where four distinct bacteria, three *Clostridia* species and *Eggerthella lenta*, each perform a critical metabolic step, although two studies have identified a positive association between dietary intake of lignan, serum enterolignan, and archaeal abundance in stool^53,54^. Due to the conversion of lignans into enterolignans capable of binding to estrogen receptors, interest in the role of lignan in human health continues to rise, with several large epidemiological studies showing the association between dietary lignan and decreased breast cancer risk^35^, coronary artery disease risk^34^, diabetes incidence and mortality^55,56^. Our data suggests that archaea may be involved in the degradation of lignan molecules into aromatic amino acid precursors in healthy people which may allow a shift of lignan metabolism away from potentially carcinogenic pathways.

Numerous well-known, large-scale multi-cohort studies published in the last several years involving tens of thousands of individuals associated with the Personalized Responses to Dietary Composition Trial (PREDICT) trial have expanded our understanding of diet-microbiota interactions, including identifying specific bacteria associated with dietary indices, food groups, and dietary molecules. In our much smaller cohort with substantially less expensive techniques and minimal sampling, we were still able to identify several significant associations between bacterial families and dietary food groups previously reported. Our success in finding known diet-microbiome relationship serves as an example of not only the robust effect of diet on the microbiota composition, but also reinforces the worthwhile contributions to our understanding of complex nature of diet-microbiome interactions that can come from both large and small studies alike.

## ACKNOWLEDGEMENTS

This study was in part funded by Bio-Rad and supported by the National Center for Advancing Translational Sciences of the National Institutes of Health under Award Number UL1TR002378. The content is solely the responsibility of the authors and does not necessarily represent the official views of the National Institutes of Health. CK is a consultant at Rebiotix/Ferring, an unpaid board member of Project Mercy and National MPS Society, in addition to serving as an associate editor at Clinical Infectious Diseases and the ASM-Health unit chair.

---

**Supplemental Figure S1.**
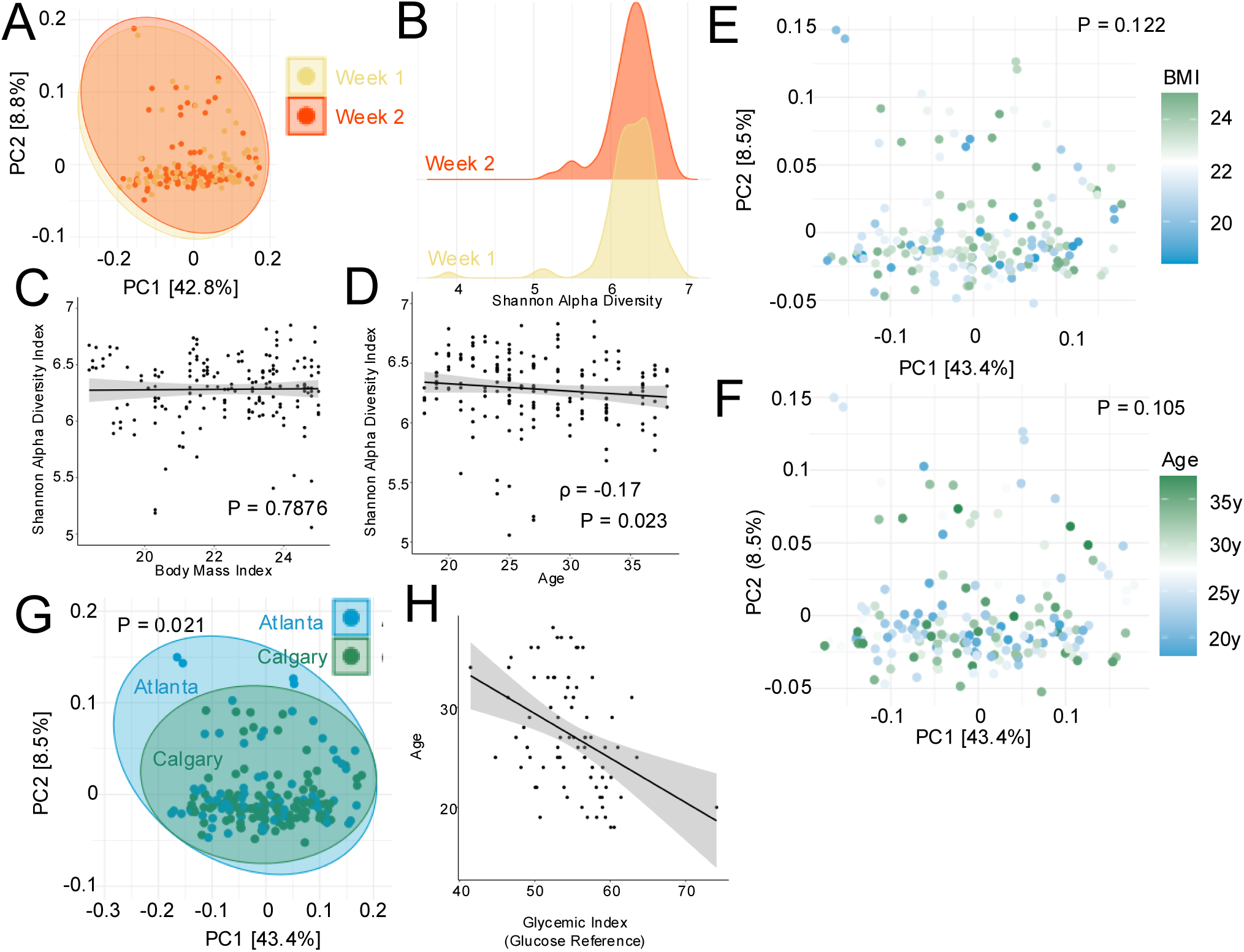
(A) Principal Component Analysis (PCoA) of 16S rRNA microbiota composition comparing samples collected from Week 1 and Week 2. (B) Distribution of Shannon Alpha Diversity of samples collected from Week 1 and Week 2. (C) Spearman correlation analysis between BMI and Shannon Alpha diversity. (D) Spearman correlation analysis between Age and Shannon Alpha Diversity index. (E) Principal Component Analysis (PCoA) of 16S rRNA microbiota composition with green-blue color shading indicating BMI. (F) Principal Component Analysis (PCoA) of 16S rRNA microbiota composition with green-blue color shading indicating age. (G) Principal Component Analysis (PCoA) of 16S rRNA microbiota composition comparing samples from Calgary and Atlanta. (H) Spearman correlation analysis between ‘Glycemic index (Glucose Reference) and Age.

## REFERENCES

1. Downer, S., Berkowitz, S. A., Harlan, T. S., Olstad, D. L. & Mozaffarian, D. Food is medicine: actions to integrate food and nutrition into healthcare. BMJ m2482 (2020) doi:10.1136/bmj.m2482.

2. Afshin, A. et al. Health effects of dietary risks in 195 countries, 1990–2017: a systematic analysis for the Global Burden of Disease Study 2017. The Lancet 393, 1958–1972 (2019).

3. Scalbert, A. et al. The food metabolome: a window over dietary exposure. Am. J. Clin. Nutr. 99, 1286–1308 (2014).

4. Ross, F. C. et al. The interplay between diet and the gut microbiome: implications for health and disease. Nat. Rev. Microbiol. 22, 671–686 (2024).

5. Sanz, Y. et al. The gut microbiome connects nutrition and human health. Nat. Rev. Gastroenterol. Hepatol. 22, 534–555 (2025).

6. Sonnenburg, E. D. & Sonnenburg, J. L. Starving our Microbial Self: The Deleterious Consequences of a Diet Deficient in Microbiota-Accessible Carbohydrates. Cell Metab. 20, 779–786 (2014).

7. Brown, E. M., Clardy, J. & Xavier, R. J. Gut microbiome lipid metabolism and its impact on host physiology. Cell Host Microbe 31, 173–186 (2023).

8. Larabi, A. B., Masson, H. L. P. & Bäumler, A. J. Bile acids as modulators of gut microbiota composition and function. Gut Microbes 15, (2023).

9. Hazleton, K. Z. et al. Dietary fat promotes antibiotic-induced Clostridioides difficile mortality in mice. NPJ Biofilms Microbiomes 8, 15 (2022).

10. Foley, M. H. et al. Bile salt hydrolases shape the bile acid landscape and restrict Clostridioides difficile growth in the murine gut. Nat. Microbiol. 8, 611–628 (2023).

11. Foley, M. H. et al. Bile salt hydrolases shape the bile acid landscape and restrict Clostridioides difficile growth in the murine gut. Nat. Microbiol. 8, 611–628 (2023).

12. Alavi, S. et al. Interpersonal Gut Microbiome Variation Drives Susceptibility and Resistance to Cholera Infection. Cell 181, 1533–1546.e13 (2020).

13. Miller, P. E. et al. Development and evaluation of a method for calculating the Healthy Eating Index-2005 using the Nutrition Data System for Research. Public Health Nutr. 14, 306–313 (2011).

14. Krebs-Smith, S. M. et al. Update of the Healthy Eating Index: HEI-2015. J. Acad. Nutr. Diet. 118, 1591–1602 (2018).

15. Godin, G. & Shephard, R. J. Leisure Time Exercise Questionnaire. PsycTESTS Dataset Preprint at 10.1037/t31334-000 (2015).

16. Casén, C. et al. Deviations in human gut microbiota: a novel diagnostic test for determining dysbiosis in patients with IBS or IBD. Aliment. Pharmacol. Ther. 42, 71–83 (2015).

17. Quast, C. et al. The SILVA ribosomal RNA gene database project: improved data processing and web-based tools. Nucleic Acids Res. 41, D590–D596 (2012).

18. Schliep, K. P. phangorn: phylogenetic analysis in R. Bioinformatics 27, 592–593 (2011).

19. Dahl, E. M., Neer, E., Bowie, K. R., Leung, E. T. & Karstens, L. microshades: An R Package for Improving Color Accessibility and Organization of Microbiome Data. Microbiol. Resour. Announc. 11, (2022).

20. Wickham, H. ggplot2. WIREs Computational Statistics 3, 180–185 (2011).

21. Douglas, G. M. et al. PICRUSt2 for prediction of metagenome functions. Nat. Biotechnol. 38, 685–688 (2020).

22. Turnbaugh, P. J. et al. The Human Microbiome Project. Nature 449, 804–810 (2007).

23. Martínez, I. et al. Gut microbiome composition is linked to whole grain-induced immunological improvements. ISME J. 7, 269–280 (2013).

24. Yang, Z. et al. Veillonella intestinal colonization promotes C. difficile infection in Crohn’s disease. Cell Host Microbe https://doi.org/10.1016/j.chom.2025.07.019 (2025) doi:10.1016/j.chom.2025.07.019.

25. Rocha, I. M. G. da et al. Pro-Inflammatory Diet Is Correlated with High Veillonella rogosae, Gut Inflammation and Clinical Relapse of Inflammatory Bowel Disease. Nutrients 15, 4148 (2023).

26. Zhang, S.-M. & Huang, S.-L. The Commensal Anaerobe Veillonella dispar Reprograms Its Lactate Metabolism and Short-Chain Fatty Acid Production during the Stationary Phase. Microbiol. Spectr. 11, (2023).

27. Miranda, P. M. et al. High salt diet exacerbates colitis in mice by decreasing Lactobacillus levels and butyrate production. Microbiome 6, 57 (2018).

28. Ferguson, J. F., et al. High dietary salt–induced DC activation underlies microbial dysbiosis-associated hypertension. JCI Insight 4, (2019).

29. Wang, C. et al. High-Salt Diet Has a Certain Impact on Protein Digestion and Gut Microbiota: A Sequencing and Proteome Combined Study. Front. Microbiol. 8, (2017).

30. Griffin, M. E., et al. *Enterococcus* peptidoglycan remodeling promotes checkpoint inhibitor cancer immunotherapy. Science (1979). 373, 1040–1046 (2021).

31. Royet, J., Gupta, D. & Dziarski, R. Peptidoglycan recognition proteins: modulators of the microbiome and inflammation. Nat. Rev. Immunol. 11, 837–851 (2011).

32. Casén, C. et al. Deviations in human gut microbiota: a novel diagnostic test for determining dysbiosis in patients with IBS or IBD. Aliment. Pharmacol. Ther. 42, 71–83 (2015).

33. Adlercreutz, H. Lignans and Human Health. Crit. Rev. Clin. Lab. Sci. 44, 483–525 (2007).

34. Hu, Y. et al. Lignan Intake and Risk of Coronary Heart Disease. J. Am. Coll. Cardiol. 78, 666–678 (2021).

35. Touillaud, M. S. et al. Dietary Lignan Intake and Postmenopausal Breast Cancer Risk by Estrogen and Progesterone Receptor Status. JNCI Journal of the National Cancer Institute 99, 475–486 (2007).

36. Asnicar, F. et al. Microbiome connections with host metabolism and habitual diet from 1,098 deeply phenotyped individuals. Nat. Med. 27, 321–332 (2021).

37. Aslam, H. et al. The effects of dairy and dairy derivatives on the gut microbiota: a systematic literature review. Gut Microbes 12, 1799533 (2020).

38. Fackelmann, G. et al. Gut microbiome signatures of vegan, vegetarian and omnivore diets and associated health outcomes across 21,561 individuals. Nat. Microbiol. 10, 41–52 (2025).

39. Kable, M. E., Chin, E. L., Huang, L., Stephensen, C. B. & Lemay, D. G. Association of Estimated Daily Lactose Consumption, Lactase Persistence Genotype (rs4988235), and Gut Microbiota in Healthy Adults in the United States. J. Nutr. 153, 2163–2173 (2023).

40. Shuai, M. et al. Multi-omics analyses reveal relationships among dairy consumption, gut microbiota and cardiometabolic health. EBioMedicine 66, 103284 (2021).

41. Aslam, H. et al. Gut Microbiome Diversity and Composition Are Associated with Habitual Dairy Intakes: A Cross-Sectional Study in Men. J. Nutr. 151, 3400–3412 (2021).

42. Castro, M., Silver, H. J., Hazleton, K., Lozupone, C. & Nicholson, M. R. The Impact of Diet on *Clostridioides difficile* Infection: A Review. J. Infect. Dis. 231, e1010–e1018 (2025).

43. Jose, S. et al. Obeticholic acid ameliorates severity of Clostridioides difficile infection in high fat diet-induced obese mice. Mucosal Immunol. 14, 500–510 (2021).

44. Hazleton, K. Z. et al. Dietary fat promotes antibiotic-induced Clostridioides difficile mortality in mice. NPJ Biofilms Microbiomes 8, 15 (2022).

45. Mefferd, C. C., et al. A High-Fat/High-Protein, Atkins-Type Diet Exacerbates *Clostridioides* (*Clostridium*) *difficile* Infection in Mice, whereas a High-Carbohydrate Diet Protects. mSystems 5, (2020).

46. Collins, M. D., et al. The Phylogeny of the Genus Clostridium: Proposal of Five New Genera and Eleven New Species Combinations. Int. J. Syst. Bacteriol. 44, 812–826 (1994).

47. Lee, J.-Y. et al. High fat intake sustains sorbitol intolerance after antibiotic-mediated Clostridia depletion from the gut microbiota. Cell 187, 1191–1205.e15 (2024).

48. Woting, A., Pfeiffer, N., Loh, G., Klaus, S. & Blaut, M. Clostridium ramosum Promotes High-Fat Diet-Induced Obesity in Gnotobiotic Mouse Models. mBio 5, (2014).

49. Bahadoran, Z. et al. Nitrate and nitrite content of vegetables, fruits, grains, legumes, dairy products, meats and processed meats. Journal of Food Composition and Analysis 51, 93– 105 (2016).

50. Jockel-Schneider, Y. et al. Nitrate-rich diet alters the composition of the oral microbiota in periodontal recall patients. J. Periodontol. 92, 1536–1545 (2021).

51. Vanhatalo, A. et al. Nitrate-responsive oral microbiome modulates nitric oxide homeostasis and blood pressure in humans. Free Radic. Biol. Med. 124, 21–30 (2018).

52. du Toit, L. et al. The Effect of Dietary Nitrate on the Oral Microbiome and Salivary Biomarkers in Individuals with High Blood Pressure. J. Nutr. 154, 2696–2706 (2024).

53. Li, Y. et al. Dietary lignans, plasma enterolactone levels, and metabolic risk in men: exploring the role of the gut microbiome. BMC Microbiol. 22, 82 (2022).

54. Lampe, J. W. et al. Colonic mucosal and exfoliome transcriptomic profiling and fecal microbiome response to a flaxseed lignan extract intervention in humans. Am. J. Clin. Nutr. 110, 377–390 (2019).

55. Wang, S. et al. Lignan Intake and Type 2 Diabetes Incidence Among US Men and Women. *JAMA Netw*. Open 7, e2426367 (2024).

56. Liu, B. et al. Lignan Intake and Mortality Among Adults with Incident Type 2 Diabetes– Prospective Cohort Studies. Am. J. Clin. Nutr. 121, 675–684 (2025).

